# Neural alignment of knowledge structures relates to human intelligence

**DOI:** 10.64898/2026.04.13.718008

**Authors:** Rebekka M. Tenderra, Stephanie Theves

## Abstract

Human general intelligence reflects stable performance correlations across diverse cognitive tasks and predicts major life outcomes. However, the relevant neural information processing mechanisms remain unclear. Structure mapping — the alignment of novel problems onto the relational structure of prior knowledge — has been proposed as a core principle in human reasoning. Combining time-resolved neural geometry analyses of task-based fMRI data with cognitive testing in a large sample, we demonstrate structural alignment of newly learned relations to preexisting knowledge representations in parietal cortex. Interindividual differences in neural alignment supported learning and reasoning and, beyond the task level, predicted the latent factor fluid intelligence. These findings provide first neural evidence that the computational principle of structure mapping contributes to individual differences in intelligence.

## Main Text

Human intelligence is psychometrically well-described: Performances across diverse cognitive tasks correlate, captured by a few broad cognitive abilities and one overarching general factor (*1*). The general factor of intelligence near-perfectly correlates with reasoning ability (*2*) and constitutes a stable trait that predicts major socioeconomic and health outcomes (*3–5*). Despite this robust psychometric characterization, human intelligence remains unspecified at the level of neural information processing. Previous research has identified correlates of intelligence in brain structure (*6*), intrinsic networks (*7*), and activation levels (*8*), implicating prefrontal-parietal cortices (*9, 10*), but the relevant processes remain unclear.

To explain psychometric observations, an intelligence-related processing mechanism should be consistent with the central role of reasoning (g_f_) in intelligence. Human reasoning often relies on structural comparison to existing knowledge representations (*11–13*). Structural abstraction and generalization via low-dimensional representations in the hippocampal system and prefrontal-parietal cortices have thus been proposed as candidate neural mechanisms (*14*). Alignment to existing knowledge structures enables inference of missing links, accelerates learning, and thereby further supports reasoning (*15, 16*) - systematic recruitment of these mechanisms may thus enhance cognitive performance more broadly (*14*). Recent work has linked intelligence to the initial formation of relational representations through integration in hippocampal cognitive maps (*17*).

Here we investigate intelligence-related differences in across-domain structure coding and generalization. Structural generalization requires a representational format that can be reused across tasks and domains, in particular through abstraction from specifics and alignment of relation vectors. These properties have been related to the hippocampal system (*18–20*) and the parietal cortex (*21, 22*). The parietal cortex processes time, space, and number within overlapping circuits (*23, 24*), representing magnitude in low-dimensional geometries that can align between tasks (*25*) suggesting relational abstraction. However, it remains unclear whether these representational mechanisms support learning and reasoning performance in a given task, much less whether interindividual differences in these mechanisms relate to general cognitive ability. An intelligence-related processing mechanism should be consistent with the central role of reasoning in intelligence, be functionally relevant for performance, predict intelligence beyond task performance, and relate to a latent factor rather than a single task measure.

To probe whether reasoning is accompanied by neural alignment to existing knowledge structures, we measured the parietal number line representation, a familiar concept that should be structurally compatible with the latent linear organization of a newly learned item ranking. We demonstrate cross-domain alignment of both structure representations in parietal cortex during inference of unseen links. Critically – consistent with the criteria above – we demonstrate that neural structural alignment supports learning and reasoning, and beyond task performance, predicts the latent factor fluid intelligence. These results identify neural structure mapping as an intelligence-related processing mechanism.

### Transitive inference performance indicates a structure representation

We probed representations of relational structure using a transitive inference paradigm (*26*) in which participants learned and reasoned over object relations on a linear-ordered dimension (Fig. 1A). Specifically, they learned the relative ordering of six nonsense objects along a latent dimension termed “flafe” through training on adjacent object pairs, then had to infer unseen rankings. In this transitive inference task, objects were presented sequentially and participants indicated for each object whether it was more or less “flafe” than the one preceding it. In a final task, participants compared Arabic digits in terms of numerical magnitude. This allowed us to assess whether newly learned object relations structurally aligned with established numerical magnitude representations. Intelligence factors, including fluid intelligence (g_f_), were assessed using a standardized test battery (*27*) on a separate day.

**Fig. 1.**
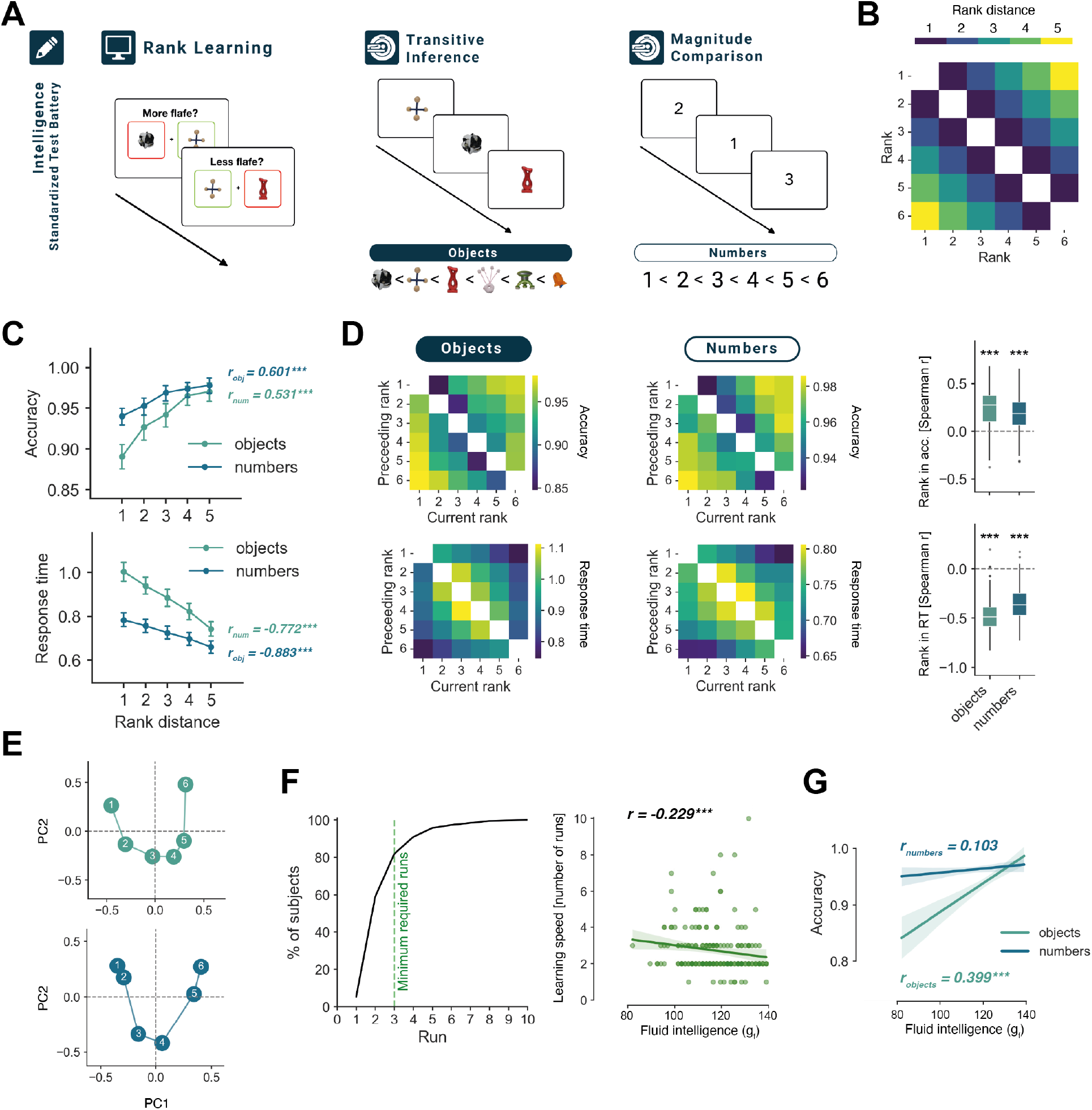
Linear structure learning relates to intelligence. **(A)** Participants completed a standardized test battery to assess fluid intelligence (g_f_). In the experimental session, they learned to compare neighboring objects on a novel dimension (“flafe”). A subsequent inference task during fMRI required rank judgements on the unseen pairs. Participants indicated for each object whether it was more or less “flafe” than the preceding one in the sequence. In a final task, they performed numerical magnitude comparisons on Arabic numbers. **(B)** Model matrix indicating rank distances between stimuli in the object transitive inference and number magnitude comparison task on a linear structure. **(C)** Accuracy increased and response times decreased with rank distance in both tasks, consistent with a line-like representation. Points denote means, error bars the 95% confidence interval. **(D)** Group average matrices of accuracy and response time (RT) per rank comparison closely follow the rank-distance model in both tasks. Boxplots show the distribution of participant-wise correlations between empirical matrices and the model. The box denotes the median and IQR, whiskers cover 1.5*IQR, points the outliers. **(E)** Classical MDS reconstruction of the average accuracy matrices of objects (top) and numbers (bottom) reveals that performance primarily reflects the factors rank-distance and distance-to-endpoint. Axes show the first and second principal component (PC). **(F)** The rank representation in reaction times in the transitive inference task (objects) correlates with intelligence (g_f_). **(G)** Most participants passed the predefined learning threshold within the first three runs (left). With increasing fluid intelligence (g_f_), participants required fewer runs to reach the criterion (right). *\*P<0*.*05; **P<0*.*01; ***P<0*.*001*.

Participants learned which of two adjacent objects was more or less “flafe” through feedback. They were trained for a minimum of 3 runs, until they reached a performance criterion to ensure knowledge of the neighboring rank pairs (final run accuracy: mean(SD) = 97.3(2.4)%). The learning rate, indicated by how many runs were required to reach criterion, correlated with fluid intelligence (Fig. 1F; r_spearman_(188) = - 0.229, P < 0.001). Performance was high overall: In the transitive inference task (accuracy: mean(SD) = 93.2(7.8)%), both for learned neighboring pairs (mean(SD) = 89.8(9.0)%) and inferred non-neighboring pairs (mean(SD) = 94.9(8.3)%), and in the subsequent number comparison task (accuracy: mean(SD) = 96.4(4.2)%). Accuracy in the inference task correlated with fluid intelligence, more than accuracy in the numerical comparison task (Fig. 1G; objects: r(188) = 0.399, P < 10^-8^; numbers: r(188) = 0.103, P = 0.158; difference: z(188) = 3.447, P < 0.001).

Both transitive inferences on the object ranks and number comparisons showed a robust symbolic distance effect (*28*): accuracy was higher and reaction time was faster, the larger the rank-distance between the two compared stimuli. This pattern indicates that inferences are supported by a representation of the linear structure, rather than by chaining local representations in which case reaction times would increase with distance. We demonstrate a symbolic distance effect when collapsing trials by rank distance (Fig. 1C; accuracy: r_obj_= 0.601, t(188) = 20.863, P < 10^-51^; r_num_= 0.531, t(188) = 17.638, P < 10^-42^; response times: r_obj_= -0.883, t(188) = -62.984, P < 10^-128^; r_num_= -0.772, t(188) = -38.299, P < 10^-91^). The effect was also evident in the correlation between pairwise accuracy and response time matrices (Fig. 1D) with the rank-distance matrix (Fig. 1B) suggesting a line-like representation of stimuli in both tasks (accuracy: r_obj_(SD)= 0.243 (0.196), t(188) = 16.988, P < 10^-40^; r_num_(SD)= 0.177 (0.181), t(188) = 13.404, P < 10^-29^; response times: r_obj_(SD)= -0.475 (0.161), t(188) = -40.474, P < 10^-94^; r_num_(SD)= -0.352 (0.168), t(188) = -28.815, P < 10^-70^). Consistent with this interpretation, the first principal component (PC) of these matrices reflected the underlying rank order. In line with previous reports in similar paradigms (*25, 29*), the second PC reflected the ranks’ distance to the endpoints, which is orthogonal to rank magnitude in this task, leading to a curved appearance of the rank representation (Fig. 1E).

### Structure representation in parietal cortex

We next examined whether line-like rank representations emerged within each task and whether they were expressed in a format that could generalize across tasks. Since we predicted that linear structure representations of numerical magnitude are reused when learning novel object ranks, we performed an ROI analysis on the Intraparietal Sulcus (IPS) where line-like number representations have previously been localized (*30, 31*). Since rank representations may arise at different latencies in the object and number tasks (*32*) and response time differed between the tasks (ΔRT(SD) = 0.172(0.177) ms, t(188) = 13.266, P < 10^-29^), we examined cross-task generalization in a time-resolved representational similarity analysis (*33, 34*). To track the emergence of line-like representations, we estimated rank-specific response patterns at each time point of the fMRI time series (TR) after stimulus onset using a finite impulse response (FIR) model and performed a rank-distance-based Representational Similarity Analysis (RSA) at each time point (Fig. 2A-B). A temporal cluster-based permutation test (P < 0.05, TFCE-corrected) identified distinct time windows after stimulus onset in which objects and numbers showed a line-like representational geometry (Fig. 2C).

**Figure 2.**
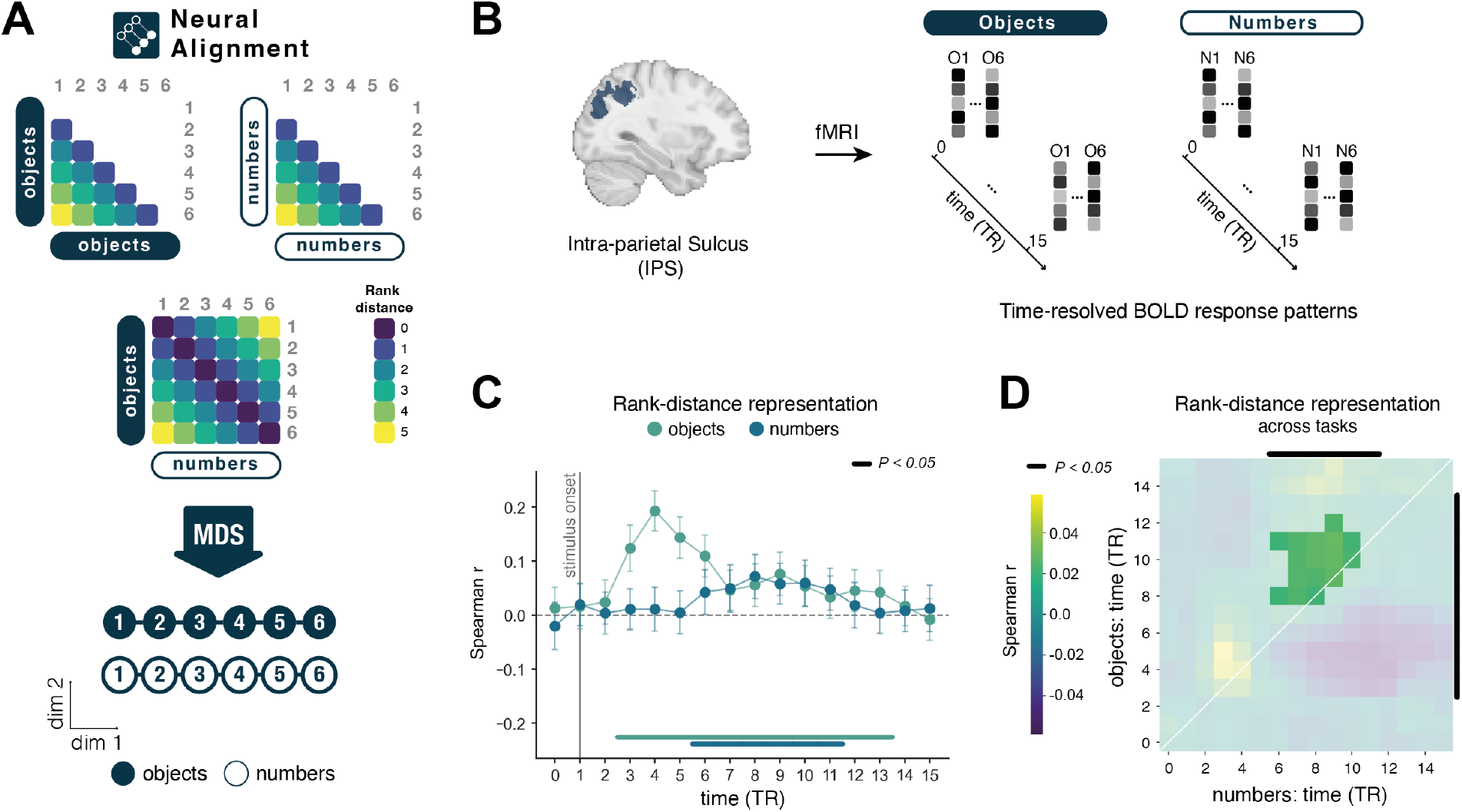
Structural alignment of object and number representations in parietal cortex. **(A)** The prediction RDM comprises rank distances reflecting a line-like arrangement of objects (object model, upper left), numbers (number model, upper right), and alignment across conditions (across model, center), illustrated as a low-dimensional embedding (bottom). **(B)** Based on fMRI data in intra-parietal sulcus (IPS), rank-specific blood oxygen level dependent (BOLD) response patterns were estimated at multiple time points after stimulus onset. **(C)** Time-resolved analysis reveals rank-distance representations emerging in IPS. Line-like representational geometry for objects (green) and numbers (blue) appeared in distinct post-stimulus time windows. Points show the mean correlation to the respective prediction RDM, error bars the 95% confidence interval. Bold lines indicate significant time windows (cluster-based permutation test, P < 0.05, TFCE-corrected). **(D)** Neural alignment occurs in a late time window. The matrix shows the mean correlation of the cross-task RDM with the prediction RDM for all time point combinations (cluster-based permutation test, P < 0.05, TFCE-corrected). Non-significant values are greyed-out. Bold black lines mark the significant time windows from (C).

### Alignment of structures across domains

To assess structural alignment between object relations in the transitive inference task and number line representations, participants’ cross-task rank Representational Dissimilarity Matrices (RDMs) were correlated with a rank-distance RDM reflecting alignment of the linear structures, overlayed by rank (cf. Fig. 2A). This was done at each pair of time points where both task-specific line-like representations had been identified. Object and number representations were structurally aligned in IPS (Fig. 2D; P < 0.05, TFCE-corrected).

### Structural alignment promotes task performance and relates to the factor fluid intelligence

The strength of structural alignment correlated positively with learning and inference performance: Participants who showed stronger structural alignment learned the object rank pairs in fewer trials (r_spearman_(188) = -0.150, P = 0.02; Fig. 3A) and responded more accurately in the inference task (r(188) = 0.166, P = 0.011), whereas the correlation between neural alignment and performance in the number comparison task (r(188) = -0.037, P = 0.692) was significantly lower (z(188) = 2.23, P = 0.013; Fig. 3B). That alignment of abstract structure representations was related to performance suggests that transitive inference in this task arises from structure mapping rather than learning item-specific values.

**Figure 3.**
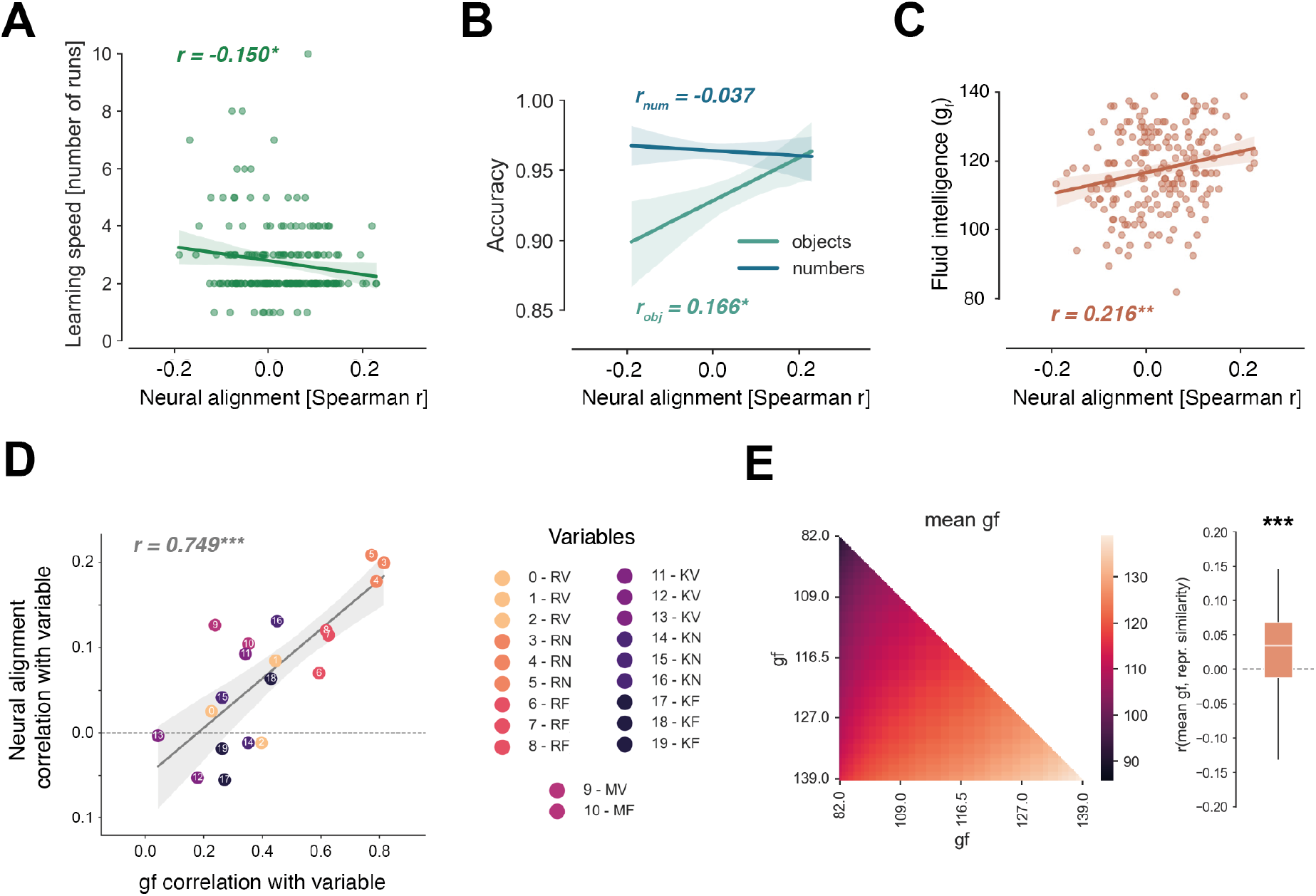
Structural alignment links to faster learning, more accurate inference, and higher intelligence. **(A)** Participants who show stronger structural alignment to numerical magnitude representations, learned the pair-wise ranking of novel objects in fewer runs. The line shows a linear model fit to the data; the shadow indicates the 95% confidence interval of the fit. **(B)** Stronger structural alignment is linked to higher accuracy in the object transitive inference task (left) but not in the number magnitude comparison task (right). The dashed grey line indicates chance level. **(C)** Structural alignment in IPS is positively related to fluid intelligence (g_f_). **(D)** Structural alignment relates to the latent factor g_f_. The higher a cognitive task loads on g_f_, the stronger it correlates with cross-task alignment in IPS. Performance variables are grouped by ability (R = reasoning, M = memory, K = knowledge) and domain (V = verbal, N = numeric, F = figural). **(E)** With decreasing g_f_, participants show more idiosyncratic cross-task alignment geometries. For each subject, we correlate pairwise mean g_f_ (this subject to all other subjects, left) with the respective pairwise representational similarity (Spearman r) of their neural cross-task alignment RDMs. Cross-participant representational similarity and g_f_ are significantly correlated. *\*P<0*.*05; **P<0*.*01; ***P<0*.*001*.

In line with our main hypothesis, structural alignment correlated positively with fluid intelligence (Fig. 3C; r(188) = 0.216, P = 0.002, controlled for nuisance covariates: r_partial_(188) = 0.209, P = 0.002). This relation persisted when controlling for task performance (r_partial_(188) = 0.167, P = 0.011), response times (r_partial_(188) = 0.207, P = 0.002), and the strength of the object and number line representations (r_partial_(188) = 0.226, P < 0.001).

If neural structural alignment relates to the latent factor fluid intelligence (g_f_), its association with cognitive performance should scale with the degree to which individual tasks load on g_f_. Consistent with this prediction, the correlation between neural alignment and performance on individual cognitive tasks increased as a function of each task’s g_f_ loading (r(20) = 0.784, P < 10^-5^, Fig. 3D).

### Representational alignment between individuals increases with g_f_

Lastly, we addressed the possibility that individuals in the lower g_f_ range align the structures consistently, yet in an alternative format. To this end, we examined whether g_f_ scaled with the between-subject similarity of cross-task representational alignment. The similarity of cross-task representational alignment decreased significantly with decreasing mean g_f_ of a participant pair (Fig. 3E; t(188) = 8.688, P < 10^-8^; controlling for nuisance covariates: t(188) = 10.108, P < 10^-10^), indicating participants in the lower g_f_ range deviated from a canonical representation rather than adopting a common alternative representation.

## Discussion

We identify neural alignment of new information to prior knowledge structures as a candidate mechanism underlying human intelligence. Using fMRI, we show that newly learned associations are mapped onto preexisting knowledge structures in parietal cortex during reasoning, and that the strength of structural alignment supports learning and inference performance and, beyond the task level, predicts the factor fluid intelligence.

We observed line-like representations of ranks in a transitive inference and number comparison task in parietal and prefrontal cortices, congruent with a previous study (*26*). Critically, we identify structural alignment of the object and number lines in parietal cortex, indicating that neural representations emerge in an abstract reusable format, which related to inference performance and scaled with fluid intelligence. This suggests that individual differences in intelligence reflect variability in representational abstraction, rather than differences in the encoding of task specifics. Abstract representational formats provide computational advantages (*18, 19*) and have been linked to faster learning, more accurate inference, and instances of zero-shot inference in animal models (*15, 35, 36*), in contexts where this format is behaviorally advantageous, such as planning (Ho et al., 2022; Theves et al., 2020). Our results suggest intelligence is related to a tendency to adopt this format.

In line with a reuse of structural knowledge in the transitive inference task, we find that neural alignment was not only related to better inference performance, but also faster rank structure learning. These findings confirm the functional relevance of neural alignment. Importantly, beyond the task level, neural structural alignment correlated positively with fluid intelligence (g_f_), which is statistically near-identical to the general factor (*2*). The correlation of neural alignment to individual cognitive tasks scaled with their respective loading on g_f_, congruent with the notion of g_f_ as a latent factor. We addressed the possibility that with decreasing g_f_, participants may align the objects and numbers in a consistent but not line-like format. Instead, we find that with decreasing g_f_ the alignment between task representations becomes more idiosyncratic. In contrast, higher g_f_ participants aligned the representations more similarly, akin to a more canonical line-like representation of magnitude.

Prior research on the neural basis of intelligence focused primarily on brain structure, functional connectivity and overall functional activation levels (*9, 10*), but neural mechanisms of how tasks are processed remained unaddressed. Together, our findings provide neural evidence that structure mapping serves as a processing principle supporting learning, reasoning performance, and more broadly intelligence (*11–14*).

Before knowledge structures can be abstracted and generalized, they need to be formed: A recent study demonstrated that the initial formation of relational representations, measured as the integration of object-locations into hippocampal-prefrontal cognitive maps, related to fluid intelligence (*17*). The hippocampus is known to be particularly engaged in new learning, whereas consolidated and schema-consistent knowledge (e.g., familiar structures such as numerical magnitude) depends on the neocortex (*15, 37, 38*). Parietal representations of abstract linear magnitude are well suited for generalization across contexts, as they appear across domains (*39*), regardless of format (*40*), and have been shown to normalize in scale (*22*). In addition to IPS—a region predominantly linked to abstract magnitude representations (*30, 31*)— a complementary searchlight analysis revealed structural generalization in dorso-medial and dorso-lateral prefrontal cortex (Fig. S1), consistent with earlier findings of magnitude representations in these regions (*26, 39–42*). In sum, the results support the idea that neural mechanisms of structural abstraction and generalization contribute to human intelligence by supporting learning, relational knowledge, and reasoning.

## Acknowledgments

We would like to thank all involved student assistants and scientific support staff for their support in data acquisition.

## Author contributions

Conceptualization: ST

Analysis: RT, ST

Investigation: RT

Visualization: RT, ST

Funding acquisition: ST

Project administration: RT, ST Supervision: ST

Writing – original draft: RT, ST

Writing – review & editing: RT, ST

## Supplementary Materials

Materials and Methods

Fig. S1

## Supplementary Materials for

## Materials and Methods

### Participants

A total of 221 adults (117 female, 99 male, 5 diverse) between the age of 18 to 35 (mean(SD)= 26.3(4.8) years) took part in the study. All participants were right-handed, had normal or corrected-to-normal vision, were native German speakers or had acquired German before the age of seven, and held a high-school diploma or an equivalent qualification. Written informed consent was obtained from all participants before participation. The study was approved by the Ethics Committee of the Medical Faculty of the University of Leipzig and carried out in accordance with the Declaration of Helsinki.

### Experimental Procedure

Participants first completed a standardized cognitive test battery. The experimental session on a separate day comprised a computerized rank learning task, and two computerized rank comparison tasks performed during fMRI acquisition.

#### Cognitive Tests

Fluid and crystallized intelligence were assessed using the I-S-T 2000 R (*27*), a commercially available German intelligence test battery. The test comprises nine reasoning subtests spanning verbal, numerical, and figural domains, as well as a general knowledge test. Fluid (g_f_) and crystallized (g_c_) intelligence scores were computed according to the test manual by transforming task-wise performance relative to a representative normative sample (healthy adults from the German general population), weighting subtests by their respective g-factor loadings, and aggregating them into normally distributed summary scores.

#### Object Transitive Inference paradigm

Participants learned the relations between six nonsense objects (“alien objects”) along a linear dimension (“flafe”, a German pseudoword), which was arbitrary and not associated with any observable stimulus feature. On each trial, two objects were presented simultaneously, accompanied by a prompt instructing participants to select the object that was either more or less “flafe”. After choosing one, colored borders around the objects indicated which response was correct (green) or incorrect (red). Feedback appeared for 1 s, either immediately upon their response or automatically after 5 s. Crucially, participants were only trained on relations between neighboring objects along the latent dimension.

The trial sequence comprised 20 unique trials (5 between-neighbor relations X 2 question types (more/less) X 2 screen positions (left-right)), which were repeated in a pseudorandomized order three times within a run. Participants completed a minimum of three learning runs and continued until they reached a predefined learning criterion at the end of a run (accuracy > 90% in the last run, and accuracy > 90% in the final repetition of unique trials).

Stimuli were selected from the NOUN database (*43*). For each participant, six objects were pseudo-randomly drawn from a pool of 26, selected to ensure high perceptual dissimilarity between stimuli based on published dissimilarity ratings.

Following learning, participants performed a transitive inference task during fMRI acquisition. This task required both neighboring and non-neighboring object comparisons, thereby probing inference beyond trained relations. On each trial, a single object was presented, and participants indicated whether the current object was more or less “flafe” than the object shown on the preceding trial.

Stimuli were presented for 0.75 s, followed by a fixation cross. Participants were instructed to respond as quickly and accurately as possible after stimulus onset. If no response was made within 2.75 s after stimulus onset, the fixation cross briefly turned blue (50 ms). The inter-trial interval was jittered according to a truncated exponential distribution (mean = 3.25 s). Response button mappings (left/right corresponding to more/less) were pseudorandomized across participants and reversed after one run.

All 30 possible object transitions were presented in pseudorandomized order and repeated three times within a run. Participants completed two runs of the task.

### Number Magnitude Comparison Task

After completion of the transitive inference task, participants performed a number comparison task Arabic digits (1-6) were presented sequentially, and participants indicated for each digit whether it was higher or lower in magnitude than the preceding digit. Presentation times and repetition of stimuli were identical to the object transitive inference test.

### Exclusion criteria

Participants were excluded from all analyses if they did not reach the learning threshold after 10 runs (n=1), their behavioral performance in learned, neighboring rank comparisons did not differ significantly from chance in any run of either of the two fMRI tasks (accuracy_mean_ < 63.3%, defined as the range containing 95% of samples from a null distribution generated from 10.000 simulated datasets with random responses, n=21), or if they missed 20% or more trials in any run (n=6). In addition, datasets were excluded from analyses if their average framewise displacement within any fMRI run exceeded 0.3 (n=14). Some participants met multiple exclusion criteria, therefore 188 participants remained for analysis.

### MRI

MRI data were acquired using a 3T Siemens MAGNETOM Skyra Connectom A scanner (Siemens, Erlangen, Germany) equipped with a 32-channel head coil. Visual stimuli were presented via a projector onto a screen viewed through a mirror mounted on the head coil. Responses were recorded using MRI-compatible button boxes.

### Anatomical imaging sequence

High-resolution T1-weighted anatomical images were acquired using a Magnetization Prepared 2 Rapid Acquisition Gradient Echo (MP2RAGE) sequence. The following acquisition parameters were used: repetition time (TR) = 5 s, echo time (TE) = 2.84 ms, inversion times (TI1, TI2) = 700 ms and 2500 ms, and flip angles = 4° (first readout) and 5° (second readout). The field of view (FOV) was 256 × 240 mm^2^ with 176 sagittal slices, providing an isotropic voxel resolution of 1 mm^3^. Full k-space sampling was performed with a partial Fourier factor of 1. To produce the final image the INV1 and INV2 images were combined, then the final T1-weighted image was generated by multiplying the intensity-normalized INV2 image with the MP2RAGE UNI image using an AFNI-based script (3dMPRAGEise, https://github.com/srikash/3dMPRAGEise). This approach enhances tissue contrast while suppressing background noise and residual artifacts.

### Functional imaging sequence

All fMRI data was recorded using a CMRR multiband echo-planar-imaging (EPI) pulse sequence (TR = 1.5s, TE = 24ms, flip angle = 70°, partial Fourier = 0.75, multiband-factor = 4, phase encoding direction = AP, voxel size = 2 mm^3^ isotropic, FOV = 204x204 mm^2^, 80 slices, interleaved slice acquisition order). The measurement volumes were angled parallel to the axis crossing the anterior and posterior commissure (ACPC). For each fMRI recording of a task, a corresponding EPI field map for estimation of the B0 field inhomogeneities was constructed based on 2 EPI reference images with opposite phase encoding direction.

### Preprocessing of MRI data

Raw DICOM files were converted into BIDS format using the dcm2bids package (version 2.1.7, https://github.com/UNFmontreal/Dcm2Bids). Results included in this manuscript come from preprocessing performed using *fMRIPrep* 22.0.1 (*44*), which is based on Nipype 1.8.4 (*45*).

#### Preprocessing of B0 inhomogeneity mappings

A *B*_*0*_-nonuniformity map (or *fieldmap*) was estimated based on two echo-planar imaging (EPI) references with topup ((*46*); FSL 6.0.5.1:57b01774).

#### Anatomical data preprocessing

The T1-weighted (T1w) image was corrected for intensity non-uniformity (INU) with N4BiasFieldCorrection (*47*), distributed with ANTs 2.3.3 ((*48*); RRID:SCR_004757), and used as T1w-reference throughout the workflow. The T1w-reference was then skull-stripped with a *Nipype* implementation of the antsBrainExtraction.sh workflow (from ANTs), using OASIS30ANTs as target template. Brain tissue segmentation of cerebrospinal fluid (CSF), white-matter (WM) and gray-matter (GM) was performed on the brain-extracted T1w using fast (FSL 6.0.5.1:57b01774, RRID:SCR_002823; (*49*)). Volume-based spatial normalization to a standard space (MNI152NLin6Asym) was performed through nonlinear registration with antsRegistration (ANTs 2.3.3), using brain-extracted versions of both T1w reference and the T1w template. The following template was selected for spatial normalization: *FSL’s MNI ICBM 152 non-linear 6th Generation Asymmetric Average Brain Stereotaxic Registration Model* [(*50*); RRID:SCR_002823; TemplateFlow ID: MNI152NLin6Asym].

#### Functional data preprocessing

For each of the 4 BOLD runs found per subject (across all tasks), the following preprocessing was performed. First, a reference volume and its skull-stripped version were generated using a custom methodology of *fMRIPrep*. Head-motion parameters with respect to the BOLD reference (transformation matrices, and six corresponding rotation and translation parameters) are estimated before any spatiotemporal filtering using mcflirt (FSL 6.0.5.1:57b01774; (*51*)). The estimated *fieldmap* was then aligned with rigid-registration to the target EPI (echo-planar imaging) reference run. The field coefficients were mapped on to the reference EPI using the transform. BOLD runs were slice-time corrected to 0.702s (0.5 of slice acquisition range 0s-1.41s) using 3dTshift from AFNI ((*52*);RRID:SCR_005927). The BOLD reference was then co-registered to the T1w reference using mri_coreg (FreeSurfer) followed by flirt ((*53*)FSL 6.0.5.1:57b01774) with the boundary-based registration (*54*) cost-function. Co-registration was configured with six degrees of freedom. Several confounding time-series were calculated based on the *preprocessed BOLD*: framewise displacement (FD), DVARS and three region-wise global signals. FD was computed using two formulations following Power (absolute sum of relative motions; (*55*)) and Jenkinson (relative root mean square displacement between affines; (*51*)). FD and DVARS are calculated for each functional run, both using their implementations in *Nipype* (following the definitions by ref. (*55*)). The three global signals are extracted within the CSF, the WM, and the whole-brain masks. The head-motion estimates calculated in the correction step were also placed within the corresponding confounds file. The confound time series derived from head motion estimates and global signals were expanded with the inclusion of temporal derivatives and quadratic terms for each (*56*). Frames that exceeded a threshold of 0.5 mm FD or 1.5 standardized DVARS were annotated as motion outliers. Additional nuisance time series are calculated by means of principal components analysis of the signal found within a thin band (*crown*) of voxels around the edge of the brain, as proposed by ref. (*57*). The BOLD time-series were resampled into standard space, generating a *preprocessed BOLD run in MNI152NLin6Asym space*. First, a reference volume and its skull-stripped version were generated using a custom methodology of *fMRIPrep*. All resamplings can be performed with *a single interpolation step* by composing all the pertinent transformations (i.e. head-motion transform matrices, susceptibility distortion correction when available, and co-registrations to anatomical and output spaces). Gridded (volumetric) resamplings were performed using antsApplyTransforms (ANTs), configured with Lanczos interpolation to minimize the smoothing effects of other kernels (*58*). Non-gridded (surface) resamplings were performed using mri_vol2surf (FreeSurfer).

Many internal operations of *fMRIPrep* use *Nilearn* 0.9.1 ((*59*);RRID:SCR_001362), mostly within the functional processing workflow. For more details of the pipeline, see the section corresponding to workflows in *fMRIPrep*’s documentation.

### Regions of Interest (ROIs)

ROIs were defined based on the Jülich Brain Atlas (*60*). Intraparietal sulcus (IPS) was defined using combined probabilistic maps of areas hIP1–hIP8 and thresholded at 50% probability.

A combined ROI of frontal (mOFC, lOFC, IFG, IFS, MFG, SFG, SFS, frontal pole, frontal gap map), parietal (SPL, IPS, IPL) and medial temporal (HC, EC, subiculum, HATA, PHC) subregions, thresholded at 50% probability, was used in the searchlight analysis.

### First-level modelling of fMRI data

First-level analyses were conducted in MNI space. Prior to modelling, data were spatially smoothed with a 3 mm FWHM Gaussian kernel.

### Quantification and statistical analysis

Associations between behavioral, neural, and intelligence measures were assessed using Pearson correlation coefficients, unless the underlying data was not continuous, in which case Spearman’s rank correlation was reported. The relation between structural generalization and cognitive performance was tested one-sided, given the directional nature of the hypothesis (cf. introduction; (*14*)) with opposite effects being of no interest, comparable to an absence of effect. Task performances relate positively to g_f_ by nature of the general factor. We note that all reported effects would be significant under two-sided testing. Whenever within-sample correlations were compared, we report Steigers’ z.

### Multi-Dimensional Scaling (MDS)

We visualized the representation of object and number relations, captured in the group-average pairwise accuracy matrix, by applying classical multidimensional scaling (MDS). The first two principal coordinates, which together captured the largest proportion of variance in the distance structure, were selected for visualization.

### Time-resolved Representational Similarity Analysis (RSA)

We estimated rank-specific responses at each TR after stimulus onset using Finite Impulse Response (FIR) models, separately for each task. This yielded one beta image per stimulus per time point per task. For each run the model additionally included six head motion and rotation parameters, framewise displacement, cosine regressors for high-pass filtering, and a constant.

Representational similarity analysis (RSA) was used to assess whether neural representations followed a line-like organization. Rank-specific activation patterns at each time point were correlated to construct representational dissimilarity matrices (RDMs) within the object task, within the number task, and across tasks. Each empirical RDM was correlated with a model RDM reflecting rank distance using Spearman’s rho. Correlation coefficients were Fisher z-transformed prior to statistical analysis.

We performed ROI-based RSA within the object transitive inference and the number magnitude comparison task to identify time windows in which line-like structure representations appeared. The resulting participant-wise data-to-model correlations for each time point were evaluated in a, one-dimensional cluster-based permutation tests (P < 0.05, TFCE-corrected, 10.000 permutations) using MNE-Python (*61*).

To assess structural generalization between tasks, rank-specific activation patterns at each time point in the object transitive inference task were correlated with rank-specific activation patterns at each time point in the number comparison task, yielding a participant-wise RDM for each pair of time points across tasks, which were then correlated with the corresponding model RDMs.

A two-dimensional temporal cluster-based permutation test (P < 0.05, TFCE-corrected, 10.000 permutations) was applied to identify contiguous time point combinations exhibiting significant rank alignment. Statistical evaluation was restricted to time point combinations for which line-like object and number line rank representations had been identified. in the one-dimensional time-resolved analysis.

This procedure identified a time window during which both object-rank and number-rank representations exhibited a line-like geometry and were aligned. To test the relationship between neural alignment to intelligence, we averaged the alignment score - defined as the correlation between empirical RDMs and the alignment model RDM across all time point combinations within the significant cluster. To test whether tasks with a higher g_f_ loading explain more variance in neural alignment we evaluated the correlation of neural alignment and each sub-task in the cognitive test battery.

While our hypothesis of generalizing prior knowledge structures (here numerical magnitude) motivated an ROI analysis of a region that represents abstract numerical magnitude, we further performed an exploratory searchlight analysis in the wider brain, again time-resolved on each time point. To determine regions showing line-like object-, number- and across-task rank representations a spatiotemporal cluster-based permutation test (P < 0.05, TFCE-corrected, 10.000 permutations) was applied to the results of the searchlight RSA. Spatial adjacency was defined by face connectivity and temporal adjacency across time points. As spatio-temporal cluster-based permutation tests in voxel-space have high computational complexity and memory demands, we tested only for concurrent line-like object-, number-, and cross-task-representations, and restricted the analysis to brain regions broadly linked to relational processing or intelligence, including grey matter voxels in medial temporal, parietal, and prefrontal cortices.

The searchlight RSA was performed with a sphere radius of 4 voxels using the rsa-toolbox (*62*).

### Representational alignment between participants

To assess whether the canonical nature of cross-task representational alignment varied with fluid intelligence, we computed pairwise similarities between participants’ cross-task alignment RDMs across all time point combinations using Spearman’s rank correlation, yielding a participant-by-participant similarity matrix. For each participant, we correlated their vector of pairwise representational similarities with the corresponding vector of pairwise mean g_f_ scores (Spearman’s rho), Fisher z-transformed the resulting coefficients, and tested them against zero using a two-sided one-sample t-test. To rule out influences of nuisance parameters, the relation was evaluated controlling for run-wise average framewise displacement (fd) and run-wise mean temporal signal-to-noise ratio (tSNR) in the selected ROI.

**Figure S1.**
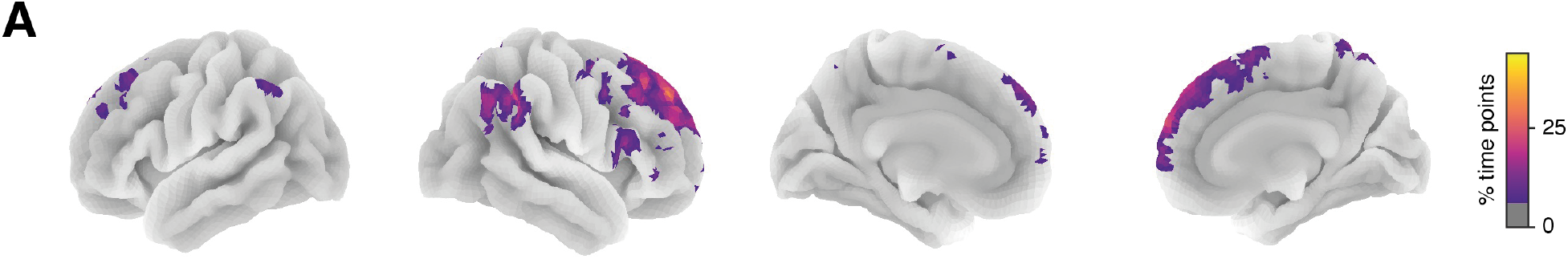
Structural generalization in dorso-medial and dorso-lateral prefrontal cortex (PFC). **(A)** The searchlight analysis of concurrent line-like object-, number, and cross-task representations (P < 0.05, TFCE-corrected, 10.000 permutations) shows structural alignment in parietal and dorso-medial frontal regions. To illustrate the spatio-temporal clusters projected to the surface, the cross-temporal mask was collapsed across time points, and therefore shows the percentage of included time points for each voxel.

